# High fidelity reconstitution of stress granules and nucleoli in mammalian cellular lysate

**DOI:** 10.1101/2020.09.14.296673

**Authors:** Brian D. Freibaum, James Messing, Peiguo Yang, Hong Joo Kim, J. Paul Taylor

## Abstract

Liquid-liquid phase separation (LLPS) is an important mechanism of intracellular organization that underlies the assembly of a variety of distinct RNP granules. Fundamental biophysical principles governing LLPS during RNP granule assembly have been revealed by simple in vitro systems consisting of several components, but these systems have limitations when studying the biology of complex, multicomponent RNP granules. Visualization of RNP granules in live cells has validated key principles revealed by simple in vitro systems, but this approach presents difficulties for interrogating biophysical features of RNP granules and provides limited ability to manipulate protein, nucleic acid, or small molecule concentrations. Here we introduce a system that builds upon recent insights into the mechanisms underlying RNP granule assembly and permits high fidelity reconstitution of stress granules and the granular component of nucleoli in mammalian cellular lysate. This system fills the gap between simple in vitro systems and live cells, and allows for a wide variety of studies of membraneless organelles.

## Introduction

A wealth of insights over the past decade has revealed the important role of biomolecular condensation in organizing cell biology through spatial and temporal control over cellular components (Banani et al., 2017). RNA metabolism in particular is governed by the formation of large, complex condensates known as RNP granules that contain hundreds of distinct proteins and RNAs (Nedelsky and Taylor, 2019). RNP granules are abundant in the nucleus (e.g., nucleoli, Cajal bodies, and speckles) and in the cytoplasm (e.g., P bodies and stress granules). Each type of RNP granule is defined by a distinct composition of proteins and RNA and plays different roles in RNA metabolism; for example, helping establish RNA three-dimensional structure, implementing splicing, conducting chemical modifications, aiding transport, regulating the translation of mRNAs, and controlling degradation. RNP granules also provide compartmentalization that has been implicated in a host of additional functions (Kedersha et al., 2013).

RNP granules assemble by liquid-liquid phase separation (LLPS), which occurs when proteins and protein-laden RNAs that are dispersed in the cytoplasm or nucleoplasm (soluble phase) coalesce into a concentrated state (condensed phase) (Brangwynne et al., 2009). In this condensed phase, the proteins and RNA behave as a single organelle with liquid-like properties, although the individual constituents remain in dynamic equilibrium with the surrounding intracellular milieu. The dynamic behavior of condensates reflects the nature of the interactions that underlie their assembly – namely, weak, transient interactions among multivalent biomolecules that form non-covalent crosslinks of varying strengths and durations. At low concentration, the individual proteins and protein-laden RNAs remain in a dispersed state. However, when the system becomes populated with sufficient crosslinks to form a system-spanning network, defined as the percolation threshold, LLPS occurs, giving rise to a condensate that is distinct from its cytoplasmic or nuclear milieu (Harmon et al., 2017). Membraneless organelles provide a strategic advantage over membrane-bound organelles because they not only concentrate macromolecules in space, but can assemble and disassemble rapidly and permit rapid exchange of constituents with the surrounding cytoplasm or nucleoplasm. The material properties of condensates, including viscosity, elasticity, and surface tension, are directly linked to condensate function, and alterations to material properties in RNP granules, and stress granules in particular, have been linked to neurodegenerative diseases such as amyotrophic lateral sclerosis (ALS) and frontotemporal dementia (FTD) (Nedelsky and Taylor, 2019).

Recent studies have yielded insight into the logic that defines the percolation threshold for complex condensates. For RNP granules, this threshold is encoded by the network of protein-protein, protein-RNA, and RNA-RNA interactions that drives LLPS (Guillen-Boixet et al., 2020; Sanders et al., 2020; Yang et al., 2020). Each component of this network contributes toward the sum of interactions required to reach the percolation threshold. Importantly, however, some components contribute more than others; specifically, the contribution of individual nodes correlates with their centrality within the network. For example, in stress granules there are ~36 proteins that, together with RNA, provide the majority of the crosslinks that set the percolation threshold beyond which LLPS occurs. Among these proteins, the most central node for the stress granule network is G3BP1, which undergoes LLPS with RNA to initiate granule assembly (Guillen-Boixet et al., 2020; Sanders et al., 2020; Yang et al., 2020). The centrality of G3BP1 in stress granule assembly has also been shown in a optogenetic system in which light-induced LLPS of G3BP1 was sufficient to drive the formation of stress granules in cells even in the absence of exogenous stress (Zhang et al., 2019). In contrast, light-induced LLPS of other stress granule proteins, including TIA1, FUS, and TDP-43, led to the formation of liquid-like droplets that had neither the biophysical properties nor the components of stress granules induced by cellular stress (Zhang et al., 2019). Correspondingly, stress granule assembly can also be triggered by raising the concentration of RNA within the core protein-RNA interaction network. This was nicely demonstrated in a recent study using yeast lysates, in which the addition of specific mRNAs was sufficient to trigger the formation of structures that closely resembled stress granules formed in intact cells (Begovich and Wilhelm, 2020).

Fundamental principles governing the assembly of RNP granules have emerged from the study of simple, reconstituted systems of one or several molecules. However, this approach has been inadequate for studying the formation of complex, multicomponent condensates such as RNP granules. Likewise, visualization of RNP granules in cells has validated many of the principles revealed by in vitro systems, but these approaches are confounded by challenges with interrogating many key features of RNP granules, most notably their material properties (e.g., viscosity, elasticity, surface tension), which are intimately related to granule function. Furthermore, such in vivo systems suffer from limited ability to rapidly and precisely manipulate protein, nucleic acid, and small molecule concentrations. Here we have taken advantage of recent insights into the mechanisms underlying RNP granule assembly to bridge the gap between highly simplistic reconstitution and intact cells. Specifically, we use insight into the network topology underlying the assembly of RNP granules to guide the methodology for their in vitro reconstitution from whole cell lysate. Thus, our manuscript describes a protein-based system in which a mammalian cellular lysate is seeded with a purified protein (i.e., G3BP1 or NPM1) to recapitulate the formation of a distinct RNP granule in a cellular lysate. Thorough analysis reveals that lysate granules induced by G3BP1 are highly similar to stress granules in their dynamic liquid properties as well as their RNA and protein composition, whereas lysate granules induced by NPM1 recapitulate the granular compartment of nucleoli.

## Results

### Seeding of purified G3BP1 in a cell lysate induces LLPS

To establish an in vitro lysate system in which LLPS and stress granule formation could be recapitulated, deeply interrogated, and experimentally manipulated, we generated a cellular extract from U2OS cells stably expressing G3BP1-GFP (Yang et al., 2020). Cells were collected by centrifugation, lysed briefly in minimal lysis buffer, centrifuged to remove cellular debris, and the supernatant was retained to create an extract that we hereafter refer to as the cellular lysate (**Fig. 1 A and Materials and Methods**). No liquid condensates or inhomogeneities were detected in these lysates by differential interference contrast (DIC) and fluorescent imaging (**Fig. 1 B**). We then added increasing concentrations of recombinant, purified G3BP1 or other proteins to this lysate while monitoring the lysate by DIC and fluorescent microscopy (**Fig. 1 A**). Dense, spherical, liquid granules containing G3BP1-GFP were induced when the concentration of G3BP1 reached a critical seeding concentration between 2.5 and 5 μM (**Fig. 1 B**), consistent with the threshold effect that characterizes LLPS. Introducing increasing amounts of the G3BP1 protein to the lysate resulted in increased condensation (**Fig. 1 B**). In the absence of lysate, purified G3BP1 alone did not form condensates with concentrations up to 100 μM (**Fig. 1 B** and data not shown), consistent with our previous findings (Yang et al., 2020) and indicating that components within the cell lysate are required for robust LLPS of G3BP1. In contrast to G3BP1, the addition of increasing concentrations of BSA or hnRNPA1 (a protein that readily undergoes LLPS) did not induce granule formation or other condensation, indicating specificity in the formation of condensates within cell lysates (**Fig. 1 C**).

**Figure 1.**
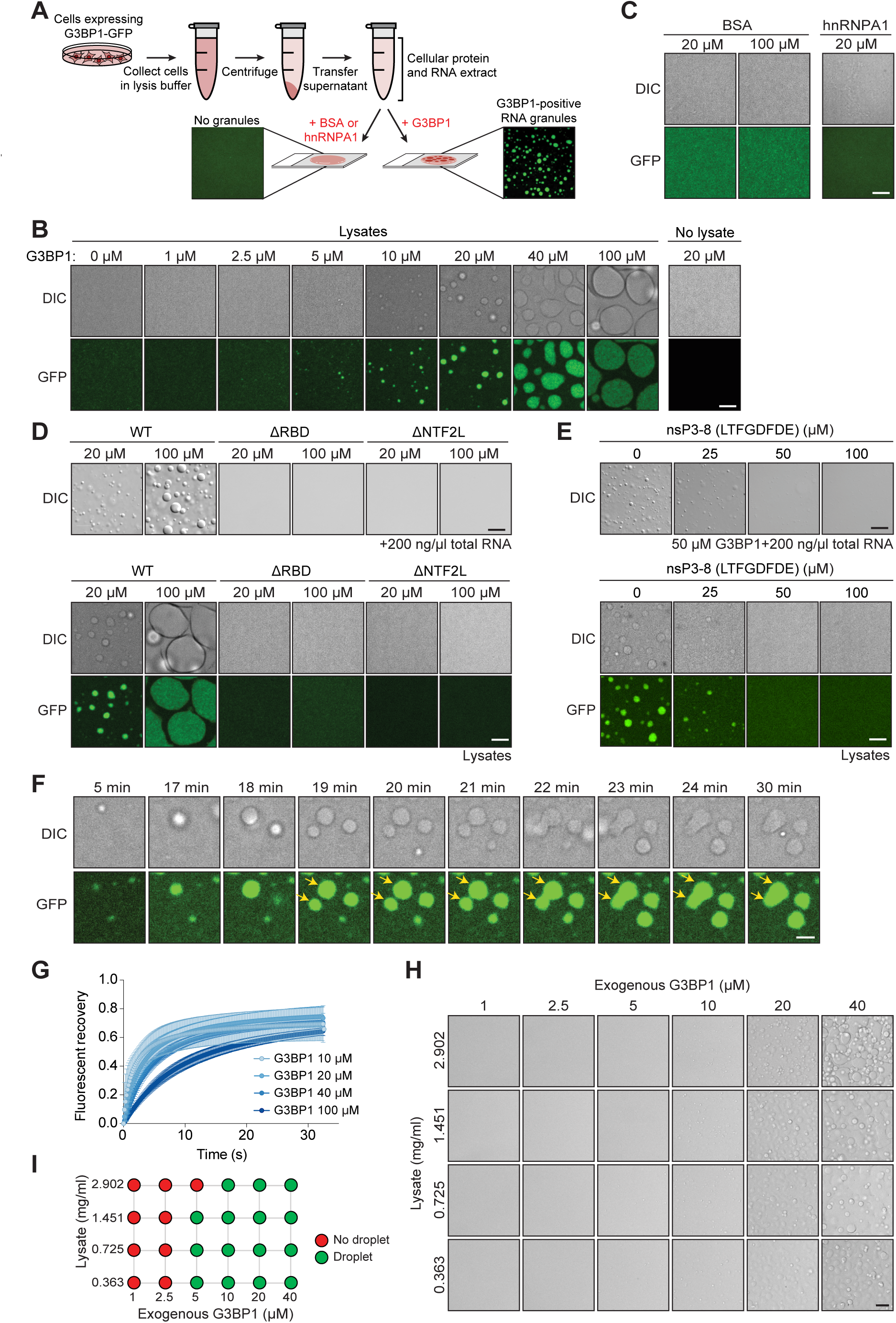
Addition of purified G3BP1 protein to a cell lysate induces LLPS to assemble G3BP1-positive granules. **(A)** Schematic illustrating the generation of lysate granules from U2OS cells stably expressing G3BP1-GFP. **(B)** Addition of G3BP1 purified protein to a cell lysate from G3BP1-GFP U2OS cells induces LLPS of G3BP1 in a dose-dependent manner when the concentration of purified G3BP1 exceeds 5 μM. LLPS is strongly inhibited in the absence of lysate. Images were taken 30 min following formation of lysate granules. Scale bar, 10 μm. **(C)** Addition of BSA or hnRNPA1 does not induce LLPS in lysates from G3BP1-GFP U2OS cells. Scale bar, 10 μm. **(D)** Purified G3BP1 lacking the RNA-binding domain (ΔRBD) or dimerization domain (ΔNTF2L) is unable to induce LLPS with RNA (top), nor does it induce LLPS in U2OS cell lysates expressing G3BP1-GFP (bottom). Scale bar, 10 μm. **(E)** Both 2-component LLPS of G3BP1 and RNA (top) as well as LLPS induced by the addition of 20 μM purified G3BP1 to U2OS cell lysates expressing G3BP1-GFP (bottom) is inhibited by the nsP3 peptide (LTFGDFDE, nsP3-8 hereafter) from Chikungunya virus. Scale bar, 10 μm. **(F)** Liquid properties of G3BP1-positive granules induced by adding 20 μM purified G3BP1 to U2OS cell lysates expressing GFP-G3BP. Granules grow and fuse over time. Scale bar, 5 μm. **(G)** FRAP performed on G3BP1-GFP within lysate granules induced by various concentrations of purified G3BP1 show that granules are mobile. Graph represents mean ± SD. **(H-I)** Brightfield phase diagram (H) with varying G3BP1 and lysate concentrations 60 min after induction (scale bar, 20 μm) and pictogram (I) indicating whether droplet formation was observed at indicated G3BP1 and lysate concentrations.

We recently showed that both the dimerization domain (NTF2L) and the RNA-binding domain of G3BP1 are required for RNA-dependent LLPS in a simple in vitro reconstitution system and also for stress granule assembly in cells (Yang et al., 2020). Consistent with these observations, G3BP1 mutants lacking these domains failed to induce LLPS of G3BP1 with RNA in vitro, as well as within our lysate system (**Fig. 1 D**), demonstrating that these G3BP1 granules have the same domain requirements for LLPS as those observed in other systems.

We next examined the effect of nsP3, a non-structural protein derived from Chikungunya virus that potently inhibits LLPS of G3BP1 by interacting with the NTF2L domain (Fros et al., 2012). LLPS of G3BP1 with RNA, as well as granule formation within our lysate system, were both inhibited in a dose-dependent manner by the addition an 8-amino acid peptide representing the FGDF motif from nsP3 (Panas et al., 2014) (**Fig. 1 E**). These results demonstrate that G3BP1-induced lysate granules reflect a G3BP1-dependent condensation that exhibits the same sensitivity to nsP3 peptide as G3BP1 LLPS with RNA in vitro and stress granule assembly in cells.

G3BP1-induced lysate granules exhibited liquid properties very similar to those of G3BP1 droplets produced in a simple in vitro reconstitution system and also stress granules assembled in cells (Yang et al., 2020), including fusion events to create granules that grow in volume over time (**Fig. 1 F and Videos 1-2**), as well as dynamic exchange of constituents with the light phase as assessed by FRAP (**Fig. 1 G and Video 3**). Furthermore, manipulating the relative concentrations of cell lysate and purified G3BP1 enabled the construction of a full phase diagram (**Fig. 1, H and I**). This diagram revealed a characteristic phase boundary, a fundamental property of LLPS (**Fig. 1, H and I**).

### Lysate granules and stress granules have similar protein composition

Stress granules are complex condensates that comprise hundreds of distinct proteins and RNAs (Khong et al., 2017; Youn et al., 2019). In cells, stress granule assembly is initiated by a rise in cytoplasmic free mRNA concentration that typically accompanies inhibition of translation initiation. The rise in mRNA concentration is sensed by G3BP1, which functions as a molecular switch to trigger stress granule assembly via RNA-dependent LLPS (Guillen-Boixet et al., 2020; Sanders et al., 2020; Yang et al., 2020). The intracellular concentration of G3BP1 sets the threshold for how high mRNA concentrations must rise to initiate stress granule assembly; indeed, with a sufficiently high concentration of G3BP1, stress granule assembly can be initiated even without a pronounced rise in cytoplasmic mRNAs. The importance of G3BP1 reflects its centrality within the core network of interactions that underlie stress granule assembly (Yang et al., 2020). We have shown this phenomenon experimentally within cells, wherein enforcing LLPS of G3BP1 using an optogenetic approach is sufficient to trigger the full cascade of stress granule assembly as assessed by protein composition of the resulting granules (Zhang et al., 2019). In contrast, enforcing LLPS of other stress granule proteins, including TIA1, FUS, and TDP-43, produced liquid droplets that did not reconstitute stress granules (Zhang et al., 2019).

We therefore hypothesized that condensation initiated by G3BP1 in cellular lysate might recapitulate full-fledged stress granule assembly. Thus, we sought to characterize the protein composition of G3BP1-induced lysate granules and assess their similarities to intracellular stress granules. We began by interrogating the protein composition of lysate granules using an indirect immunofluorescence approach. To visualize the incorporation of endogenous proteins into lysate granules, we pre-conjugated primary antibodies with secondary antibody and added these to the lysate granules immediately following induction with 20 μM G3BP1. We used antibodies against six proteins known to be stress granule constituents (CAPRIN1, PABPC1, PRRC2C, USP10, UBAP2L, CSDE1) and all six were confirmed to accumulate in lysate granules (**Fig. 2 A**). By contrast, when we used antibodies against a collection of proteins known to be absent from stress granules (actin, DCP1A, FKBP12, TOM20, GOLGA3) all were found to be absent from lysate granules (**Fig. 2 A**). To further assess lysate granule composition, we used CRISPR-Cas9 in U2OS cells to introduce fluorescent tags into endogenous genes. For example, we created dual-tagged cell U2OS lines in which endogenous G3BP1 and TDP-43 were fused to mRuby3 and GFP, respectively. When lysates were generated from these cells and supplemented with purified G3BP1, these endogenous proteins were robustly recruited to lysate granules (**Fig. 2 B**). Similarly, we also introduced fluorescent tags into the genes encoding ATXN2L and TIA1. These canonical stress granule constituents were also robustly recruited to lysate granules (**Fig. 2 B**). We also made use of the observation that overexpression of numerous stress granule constituent proteins is known to promote stress granule assembly (Kedersha et al., 2016). This phenomenon reflects positive cooperativity, which lowers the network-encoded threshold for LLPS (Yang et al., 2020). Consistent with prior observations in cells, we found that lysate generated from cells overexpressing FUS, hnRNPA1, or caprin 1 strongly promoted lysate granule assembly (**Fig. 2 C**).

**Figure 2.**
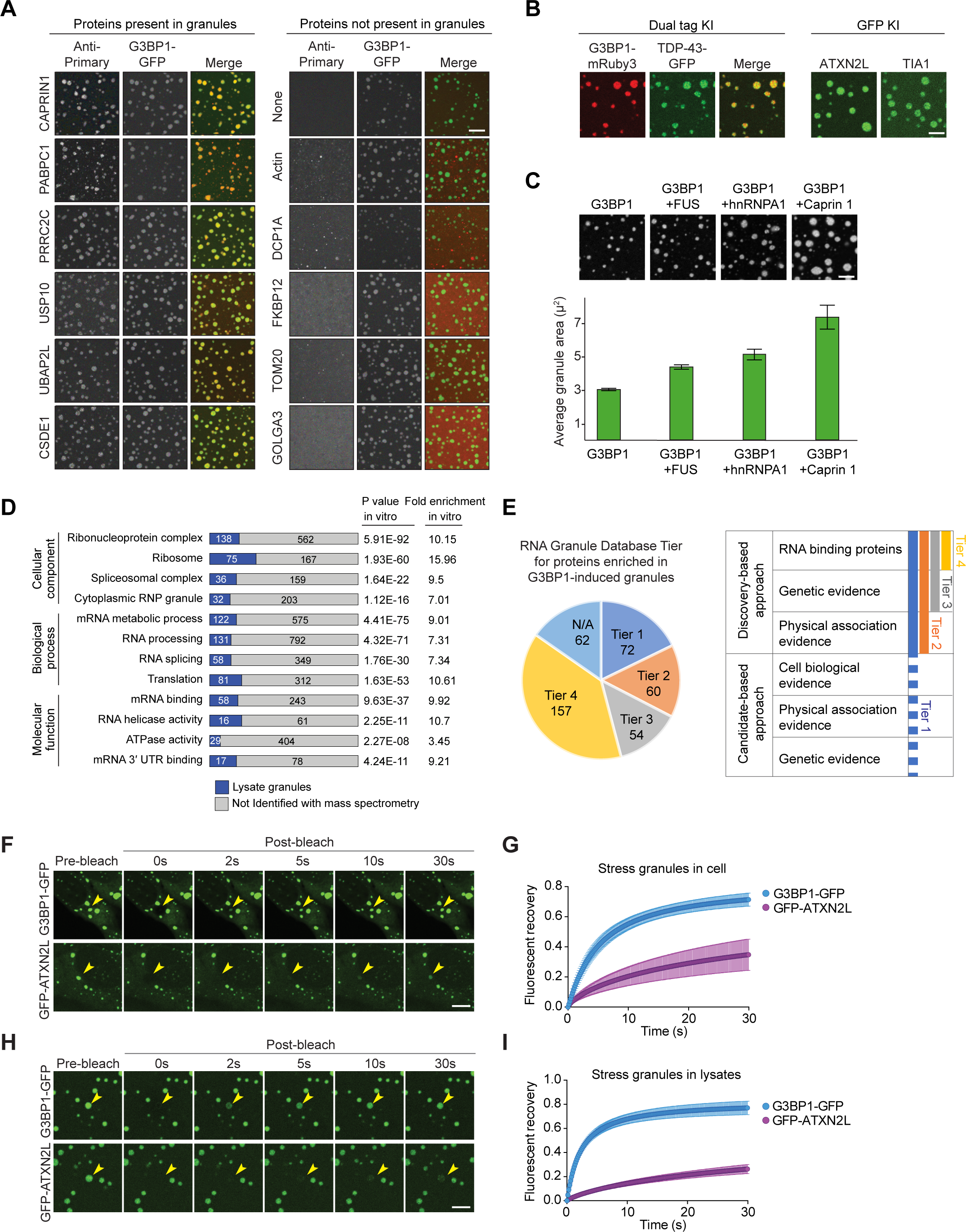
Protein composition of G3BP1-induced lysate granules resembles the protein composition of stress granules in cells. **(A)** Primary antibodies conjugated to Alexa Fluor 647 secondary antibody were used to visualize proteins known to be stress granule constituents (left) and proteins known to be absent from stress granules (right). Scale bar, 10 μm. **(B)** Endogenous canonical stress granule constituents (TDP-43, ATXN2L, TIA1) were fluorescently tagged using CRISPR-Cas9 knock-in in U2OS cells. Endogenous TDP-43, ATXN2L, and TIA1 all localized to lysate granules. Scale bar, 10 μm. **(C)** Overexpression of stress granule constituents (FUS, hnRNPA1, or caprin 1) by transfection resulted in larger granules, demonstrating positive cooperativity. Lysates were seeded at equal concentrations. Images show granules 2 h after induction with 20 μM G3BP1 and associated quantification of average granule size at 2 h. Scale bar, 10 μm. Graph represents mean ± SD. **(D)** Gene ontology analysis of proteins enriched within G3BP1-induced lysate granules reveals that lysate granules share common features with cellular RNA granules. **(E)** Most proteins identified within lysate granules appear within the RNP granule database (http://rnagranuledb.lunenfeld.ca/) as Tier 1-4 proteins, indicating that lysate granules share common components with stress granules. **(F-I)** FRAP performed on G3BP1-GFP and GFP-ATXN2L shows similar mobility between stress granules within U2OS cells induced with 0.5 mM NaAsO_2_ (F,G) and lysate granules induced with 20 μM purified G3BP1 (H,I). All images were acquired 30 min after induction. Scale bar, 10 μm. Graphs in (G) and (I) represent mean ± SD.

We also performed a comprehensive analysis of proteins recruited to G3BP1-induced lysate granules using mass spectrometry. For these experiments, we used LC/MS/MS to analyze sedimented granules induced by 20 μM purified G3BP1, using an unseeded lysate as a negative control. We found 405 proteins enriched in G3BP1-induced granules compared to control, including 267 proteins enriched >10 fold relative to control (**Table S1**). Gene ontology analysis revealed that the lysate granule proteome was highly enriched for proteins known to comprise RNP complexes, stress granules, and other aspects of RNA metabolism (**Fig. 2 D**). To assess the similarity of this proteome to the proteome of stress granules assembled in cells, we compared the proteins found in lysate granules to the list of stress granule proteins catalogued in the RNA Granule Database, an online catalog of proteins that have been reported in the literature as constituents of stress granules (Youn, submitted). Proteins within this curated database are assigned to “quality tiers” based on the extent of evidence and validation identifying each protein as a constituent of stress granules, with Tier 1 proteins representing the most extensively validated proteins. Of the 405 total proteins identified in the lysate granules, 343 were present in the RNA granule database (**Fig. 2 E**). This comparison indicates that the protein composition of G3BP1-induced lysate granules represents a high-fidelity recapitulation of stress granules that arise within cells.

Like other complex biological condensates, stress granules are highly dynamic structures in which constituents exchange between the dense phase and the surrounding cellular milieu with differing mobilities. For example, FRAP analysis demonstrated that both G3BP1 and ATXN2L are mobile, although G3BP1 shows a more rapid recovery and larger mobile fraction than ATXN2L after photobleaching (**Fig. 2, F and G and Video 4**). Induction of lysate granules with 20 μM seed G3BP1 produced granules similar in size (1-3 μm) to those induced in cells by arsenite treatment and, remarkably, FRAP analysis demonstrated that the relative dynamics of G3BP1 and ATXN2L were preserved in lysate granules (**Fig. 2, H and I and Video 5**).

Taken together, these experiments demonstrate that lysate granules are condensates that arise through LLPS triggered by G3BP1 specifically. They follow the molecular rules of G3BP1 LLPS as defined in previous studies, and are complex, dynamic condensates that recapitulate the protein constituents of stress granules with high fidelity.

### Lysate granules and stress granules have highly similar RNA composition

Stress granules are composed not only of proteins but also mRNAs. The specific enrichment of mRNAs in stress granules is consistent with the binding preference of G3BP1 and other stress granule proteins for mRNA (Yang et al., 2020). While essentially every mRNA, and some noncoding RNAs, are targeted to stress granules, the targeting efficiency varies widely, with relative enrichment of some transcripts and relative depletion of others (Khong et al., 2017). mRNA accumulation in stress granules correlates with longer coding and UTR regions as well as poor translatability. We therefore next investigated the RNA composition of lysate granules to assess similarity to stress granules assembled in cells. We began by examining the presence of mRNA within lysate granules using a fluorescent oligo-dT probe, which revealed enrichment of mRNA within granules (**Fig. 3 A**). Indeed, the RNA component seems essential to lysate granule assembly since the addition of RNase to the lysate prevented assembly of granules (**Fig. 3 A**).

**Figure 3.**
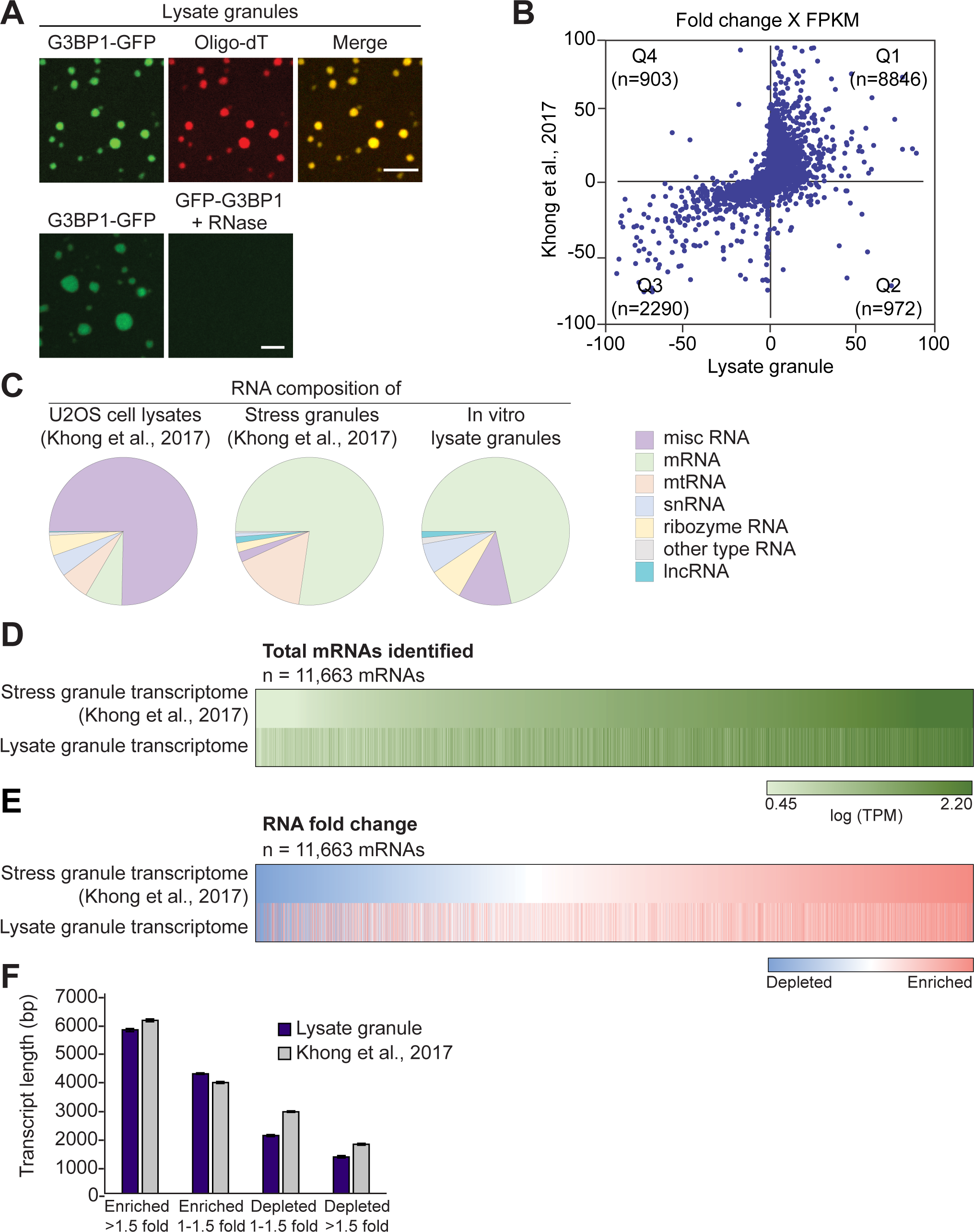
The RNA composition of G3BP1-induced lysate granules resembles the RNA composition of stress granules in cells. **(A)** Granules induced with 20 μM G3BP1 incorporated labeled oligo-dT RNA (top). The addition of RNase (1 mg/ml) prevented the formation of granules (bottom). Images were obtained 30 min following induction with G3BP1. Scale bar, 10 μm. **(B)** Relationship between fold change (stress granule vs lysate granule) in fragments per kilobase million (FPKM) of RNAs in G3BP1-induced lysate granules (X axis) compared to a published stress granule dataset (Khong et al., 2017) (Y axis). **(C)** RNA composition of whole cell lysates (left), stress granules (middle), and lysate granules (right) analyzed by RNA-seq. Both stress granules and G3BP1-induced lysate granules primarily contain mRNA compared to the lysate from which they were derived. The RNA composition of G3BP1-induced lysate granules is highly similar to the published stress granule transcriptome (Khong et al., 2017). **(D)** Individual mRNAs identified in both the stress granule transcriptome (Khong et al., 2017) and G3BP1-induced lysate granules were color-coded from the most prevalent to the least prevalent within the granule. Log(TPM) (transcripts per million) was used for color coding such that mRNAs with log(TPM) ≤ 0.45 were assigned the lightest shade of green whereas mRNAs with log(TPM) ≥ 2.20 were assigned the darkest shade of green. **(E)** mRNAs that were identified in both the stress granule transcriptome (Khong et al., 2017) and G3BP1-induced lysate granules were rank ordered from the most depleted (blue) to the most enriched (red) within the granule as compared with their respective lysates. **(F)** Average mRNA transcript length is positively corelated with the level of granule enrichment or depletion in both stress granules and lysate granules. Graph represents mean ± SEM.

We next performed RNA-seq to comprehensively assess the RNA composition of lysate granules, using RNA isolated from lysates prior to induction with G3BP1 as a control. We compared these datasets with the RNA-seq results from a previously published study of the stress granule transcriptome (Khong et al., 2017) (**Table S2**). We found strong similarity in the RNA transcriptomes of lysate granules and stress granules assembled in cells (**Fig. 3 B and Table S3**). Moreover, we found strong similarity in the classes of RNAs enriched in lysate granules and stress granules assembled in cells. Specifically, the transcriptomes of both lysate granules and stress granules were highly enriched in mRNAs and to a lesser extent various classes of non-coding RNAs such as lncRNAs and pseudogene RNAs (**Fig. 3 C and Table S3**). One notable difference was an enrichment for mitochondrial RNAs in stress granules assembled in cells that was not present in lysate granules (**Fig. 3 C and Table S3**). This could represent a true difference in composition between lysate granules and stress granules assembled in cells; alternatively, it is possible that this reflects co-purification of mitochondria with stress granules during isolation from intact cells. The lysate granules also contained a greater quantity of miscRNA and ribozyme RNA than stress granules, although these RNAs were still strongly depleted relative to control, as also observed with stress granules assembled in cells (**Fig. 3 C and Table S3**).

Our RNA-seq analysis identified 11,681 specific mRNAs within lysate granules, of which nearly all (11,663, 99.8%) were also found in the stress granule transcriptome (Khong et al., 2017). To visualize the similarity between these two datasets at the level of individual mRNAs, we generated a heat map that rank ordered mRNAs from the least prevalent to the most prevalent within the stress granule proteome, revealing a strong correlation between the mRNA composition of stress granules and lysate granules (**Fig. 3 D**). When these same mRNAs were analyzed according to depletion or enrichment in granules as compared to the respective lysates from which they were derived, the heat maps were strikingly similar (**Fig. 3 E**).

Previous analysis of the stress granule transcriptome (Khong et al., 2017) and studies of LLPS driven by G3BP1 and RNA (Yang et al., 2020) have also revealed that RNA length is an important feature of RNA that drives LLPS and recruitment into stress granules. Indeed, we found a strong similarity in the relationship between depletion/enrichment and transcript length when we compared the transcriptome of stress granules and lysate granules (**Fig. 3 F**). Altogether, these analyses demonstrate that granule assembly leads to enrichment in the same classes of RNA, as well as RNA with similar features, irrespective of whether granules are seeded by G3BP1 in lysate or assembled in response to stress in cells. Thus, lysate granules faithfully recapitulate the RNA composition of stress granules assembled in cells.

### Addition of exogenous NPM1 induces a distinct lysate granule

G3BP1 is the protein of highest centrality within the stress granule network and as such plays an outsized role in establishing the composition, and therefore identity, of stress granules (Guillen-Boixet et al., 2020; Sanders et al., 2020; Yang et al., 2020; Zhang et al., 2019). Other proteins have been implicated as perhaps playing a similar critical role in the assembly of other biomolecular condensates. For example, NPM1 is an RNA-binding protein that occupies a central position in the network of interactions that underlie formation of the liquid, granular component of the nucleolus (Mitrea et al., 2016). Indeed, analogous to G3BP1, NPM1 oligomers undergo LLPS when mixed with nucleolar proteins and rRNAs (Mitrea et al., 2016). Thus, we hypothesized that recombinant, purified NPM1 might induce complex biomolecular condensation in cellular lysate to form the liquid, granular component of the nucleolus.

Similar to our observations using purified G3BP1, the addition of recombinant, purified NPM1 to lysate resulted in concentration-dependent condensation above a threshold concentration of approximately 5 μM (**Fig. 4 A**). When examined by FRAP, the dynamic exchange of NPM1 in these lysate granules closely matched the rate of NPM1 dynamic exchange in nucleoli of intact cells (**Fig. 4, B and C, and Video 6**). Importantly, these NPM1-induced granules are distinct from G3BP1-induced granules. For example, in contrast to the potent inhibition of G3BP1-induced granules by the addition of viral nsP3 peptide, the formation of NPM1-induced granules was unaffected by the addition of nsP3 (**Fig. 4 D**), demonstrating that assembly of NPM1-induced granules does not require LLPS of G3BP1.

**Figure 4.**
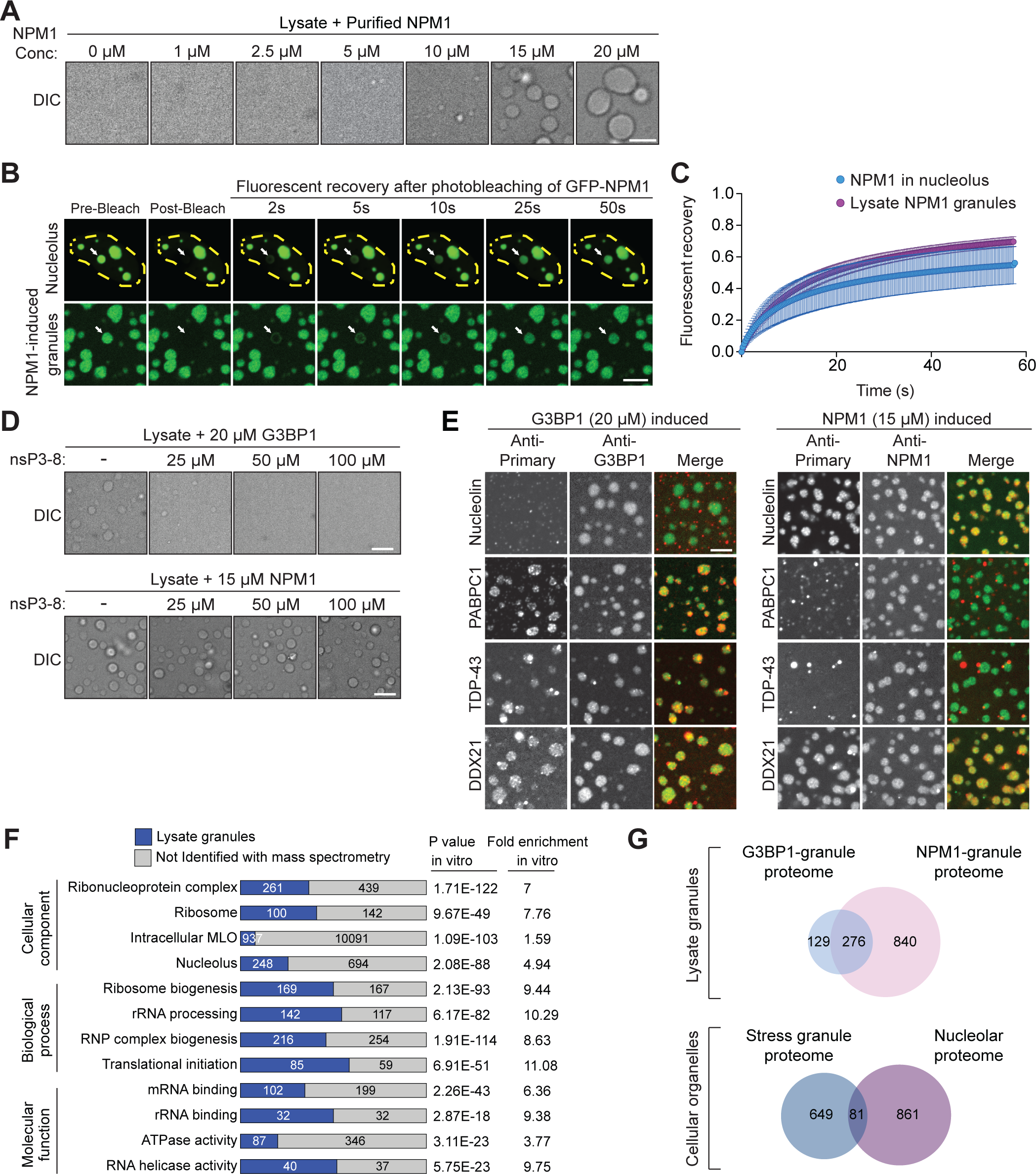
Addition of NPM1 to a cell lysate induces a distinct lysate granule that recapitulates the composition and properties of the nucleolus. **(A)** Addition of purified NPM1 protein to a cell lysate induces LLPS in a dose-dependent manner when the concentration of NPM1 exceeds 5 μM. Scale bar, 5 μm. **(B**,**C)** FRAP performed on cells or lysate from cells transiently expressing GFP-NPM1 shows similar mobility between the nucleolar NPM1 in intact U2OS cells and NPM1 in lysate granules induced with 15 μM purified NPM1. All images were acquired 30 min after induction. Scale bar, 10 μm. Graph represents mean ± standard deviation. **(D)** The nsP3-8 peptide from Chikungunya virus strongly inhibits the formation of G3BP1-induced lysate granules (top) but has no effect on the formation of NPM1-induced lysate granules (bottom). Scale bars, 10 μm. **(E)** Indicated primary antibodies were conjugated with Alexa Fluor 647 secondary, and either anti-G3BP1 or anti-NPM1 were conjugated with Alexa Fluor 488 secondary. Conjugated antibodies were added to U2OS lysates along with purified G3BP1 or NPM1. Scale bar, 10 μm. **(F)** Gene ontology analysis of proteins enriched within NPM1-induced (15 μM) lysate granules reveals that lysate granules show nucleolar features as well as some common features with G3BP1-induced (20 μM) lysate granules. **(G)** Venn diagrams showing the overlap of proteins identified through mass spectrometry between NPM1-induced lysate granules (15 μM) and G3BP1-induced lysate granules (20 μM), as well as the overlap of proteins previously identified in stress granules (RNA Granule Database, see main text) and the nucleolar proteome as assessed by GO analysis of the PANTHER database.

Next, we sought to characterize the protein composition of NPM1-induced lysate granules and compare this to the composition of nucleoli in cells as well as G3BP1-induced lysate granules. As before, we began by interrogating the protein composition of lysate granules using an indirect immunofluorescence approach. Remarkably, nucleolin, a marker enriched in nucleoli but not stress granules, was strongly recruited to NPM1-induced granules but not G3BP1-induced granules (**Fig. 4 E**). In contrast, PABPC1 and TDP-43, which are found in stress granules but not nucleoli, were strongly recruited to G3BP1-induced granules but not NPM1-induced granules (**Fig. 4 E**). Finally, DDX21, which is a constituent of both stress granules and nucleoli, was found in lysate granules induced by either G3BP1 or NPM1 (**Fig. 4 E**).

We next performed a comprehensive analysis of proteins recruited to NPM1-induced granules using mass spectrometry (**Table S4**). Remarkably, gene ontology analysis of the NPM1-induced granule proteome revealed strong enrichment of nucleolar components (**Fig. 4 F**). Notably, the proteome of NPM1-induced granules shared some features of the G3BP1-induced granule proteome, including enrichment of proteins with ATPase activity, helicase activity, and RNP components, which are typical constituents of RNP granules in cells (**Fig. 4 F**). This overlap was reflected in 276 specific proteins that were found in the proteomes of both NPM1-induced granules and G3BP1-induced granules (**Fig. 4 G**), a compositional overlap that mirrors that observed between stress granules and nucleoli in cells. We also noted that the NPM1-induced lysate granule proteome was substantially larger than that of the G3BP1-induced lysate granule proteome, again mirroring the relative complexities of the nucleolus versus stress granules in cells (**Fig. 4 G**).

## Discussion

This study describes faithful recapitulation of two different types of RNP granules within a highly tractable cell lysate system. This system is a valuable tool for studying the biology of RNP granules and bridges the gap between live cells and simple in vitro systems consisting of just a few purified components. This tool also provides novel insight into the mechanism whereby condensates establish and maintain distinct compositions. Indeed, the central observation that stress granules were reconstituted within cellular lysate by increasing the concentration of a single protein, G3BP1, fulfilled several predictions that arose from earlier work. It has been previously suggested that cells, and in particular dynamic structures such as RNP granules, are self-organizing systems whose principles of organization are determined by the intrinsic properties of the components of each structure (Misteli, 2001; Misteli, 2008). Until recently, however, there have been few ways in which this concept has been experimentally testable.

RNP granules arise by LLPS. We now appreciate that the percolation threshold for LLPS in RNP granule assembly is encoded by the network of weak, transient protein-protein, protein-RNA, and RNA-RNA interactions (Guillen-Boixet et al., 2020; Sanders et al., 2020; Yang et al., 2020). Each component of this network contributes toward the sum of interactions necessary to breach the percolation threshold. Importantly, however, some components contribute more than others. The contribution of individual proteins to stress granule assembly has been assessed by unbiased, whole-genome genetic screening (Yang et al., 2020). Moreover, the network of interactions underlying stress granule assembly in U2OS cells has now been elucidated based on stress granule proteomics (Jain et al., 2016; Markmiller et al., 2018; Youn et al., 2019).

Integration of the genetic screen results with the network of interactions has found a correlation between a protein’s centrality within the stress granule network and its importance in stress granule assembly (Yang et al., 2020). On this basis, it has been suggested that one could manipulate individual stress granule proteins with predictable outcomes (Yang et al., 2020). One illustration of this concept is the well-appreciated phenomenon whereby overexpression of some stress granule proteins in cells is sufficient to trigger condensation in the absence of exogenous stress (Kedersha et al., 2016).

The most central nodes in the stress granule interaction network – predicted to lend the greatest contribution to stress granules assembly – are G3BP1 and G3BP2 (Yang et al., 2020). Consistent with this finding was the earlier observation that double knockout of G3BP1 and G3BP2 prevents stress granule assembly in response to arsenite (Kedersha et al., 2016). The importance of G3BP1 in stress granule assembly has also been shown in an optogenetic system in which light-induced LLPS of G3BP1 in cells was sufficient to drive the formation of stress granules even in the absence of exogenous stress (Zhang et al., 2019). This is not a feature of any stress granule protein, since light-induced LLPS of TIA1, FUS, or TDP-43 does not trigger stress granule assembly (Zhang et al., 2019). Thus, the ability to trigger stress granule assembly in lysate by supplementation with purified G3BP1 fulfills a prediction built on a wealth of recent in vitro and cell-based studies. Moreover, the likely generalizability of this phenomenon is illustrated by the finding that increasing the concentration of NPM1 created a distinct condensate that recapitulates features of the nucleolar granular component, consistent with the central role of NPM1 in the network of interactions constituting this distinct condensate in cells. These findings are also consistent with a recent study using yeast lysate, in which the addition of specific mRNAs was sufficient to trigger the formation of structures that feature characteristics of stress granules (Begovich and Wilhelm, 2020). The extent to which supplementation of lysates with RNAs can fully recapitulate stress granules or other RNP granule types awaits in-depth analysis of composition through methods such as proteomics and RNA-seq.

Beyond serving as a dramatic illustration of the self-organizing principles that exist inside cells, this study also provides a useful tool for studying biomolecular condensation in general. Indeed, this system allows investigators to exploit insights into the logic that underlies the network-encoded percolation threshold for many different types of condensates. With foreknowledge of the network underlying a condensate, as well as the centrality of each node within the network, one can predict how to construct an individual condensate within a lysate; namely, by using the central constituent to breach the percolation threshold and trigger the formation of a condensate with a specific composition. Within this lysate system, it is possible to identify causal relationships between experimental manipulations and changes in granule properties independent of potentially confounding signaling pathways occurring within cells. These manipulations could include fine-tuning specific properties (e.g., salt, pH) of the milieu in which the condensates form, or genetically modifying specific constituents by introducing mutations. These mutations could be designed to interrogate the function of a specific constituent, or, importantly, could replicate a disease context. Indeed, understanding precisely how disease-causing mutations lead to changes in material properties of specific condensates remains one of the most vital unanswered questions in the field of neurodegeneration and ALS/FTD in particular.

We propose that this system could be used in studies of aging (e.g., modeling maturation of LLPS-dependent structures) and is highly amenable to screening for drugs, metabolites, or genes that impact LLPS-induced condensates. Furthermore, lysate granules settle on the surface of imaging chambers in a manner that could allow examination of granule substructures using advanced imaging techniques, including super-resolution microscopy.

Moreover, this system is ideal for mechanistic studies in which precisely manipulating the concentrations of individual constituents can enable deeper understanding of the relationship between complex networks and condensate properties.

## Materials and Methods

### Cell culture and transfection

U2OS cells were purchased from ATCC. Cells were cultured in Dulbecco’s modified Eagle’s medium (HyClone) supplemented with 10% fetal bovine serum (HyClone SH30071.03 and SH30396.03) and maintained at 37°C in a humidified incubator with 5% CO_2_. Lipofectamine 2000 (Thermo Fisher Scientific, 11668019) for U2OS cells were used for transient transfections according to the manufacturer’s instructions. U2OS cells stably expressing G3BP1-GFP have been previously described (Figley et al., 2014).

### Generation of knock-in cell lines

Genetically modified U2OS cells were generated using CRISPR-Cas9 technology. Briefly, 400,000 U2OS cells were transiently co-transfected with 200 ng gRNA expression plasmid (cloned into Addgene 43860), 500 ng Cas9 expression plasmid (Addgene 43945), 200 ng pMaxGFP, and 500 ng donor plasmid via nucleofection (Lonza, 4D-Nucleofector X-unit) using solution P3, program CM-104 in small cuvettes according to the manufacturer’s recommended protocol. Cells were single-cell sorted by FACS to enrich for GFP+ (transfected) cells, clonally selected, and verified for the desired targeted modification via PCR-based assays and targeted deep sequencing.

### Preparation for lysis

To prepare cells for use in in vitro lysate LLPS, cells were grown to confluency in 10-cm cell culture treated dishes (Corning). To harvest cells, media was aspirated and cells were washed with PBS. 5 mL PBS was added following the wash and cells were scraped to detach. After detachment, cells were centrifuged at 500 ×*g* for 3 min. PBS was removed by aspiration leaving the cell pellet untouched. Cell pellets were stored at −80°C for up to 3 months.

### Protein purification

Recombinant G3BP1 WT and mutant proteins were purified as described previously (Yang et al., 2020). hnRNPA1 was purified as described previously (Molliex et al., 2015). Purified recombinant NPM1 was received as a gift from the Kriwacki Lab (St. Jude Children’s Research Hospital, SJCRH). BSA was purchased from Sigma.

### Peptide synthesis

The Nsp3_8 WT peptide (LTFGDFDE) was synthesized by the Hartwell Center for Bioinformatics and Biotechnology at SJCRH, Molecular Synthesis Resource, using standard solid phase peptide synthesis chemistry. The peptide was reconstituted from lyophilized form into DMSO or lysis buffer (ddH_2_O, 50 mM Tris-HCl (pH 7.0), 0.5% NP-40) for experimental use with the 2-component LLPS or lysate systems respectively.

### Liquid-liquid phase separation of purified G3BP1 and RNA

LLPS of purified G3BP1 WT or mutant at the indicated concentration was induced by addition of 200 ng/mL RNA. The samples were mixed in low binding tubes (COSTAR 3206) and transferred to a sandwiched chamber created by cover glass and a glass slide with a double-sided spacer (Sigma GBL654002). Samples were observed by DIC using a Leica DMi8 microscope with 20× objective. All images were captured within 5 min after LLPS induction. The buffer for LLPS contained 150 mM NaCl and 50 mM HEPES pH 7.5. Total RNA was isolated from U2OS cells using TRIzol (Thermo Fisher Scientific) and the concentration of RNA was measured by NanoDrop (Thermo Fisher Scientific). For Nsp3_8 peptide experiments, the peptide was mixed with G3BP1 prior to the addition of RNA.

### Cell lysis and induction of LLPS from lysate

Cell pellets were removed from −80°C and thawed at room temperature for 2 min. 250 μl lysis buffer (ddH_2_O, 50 mM Tris-HCl (pH 7.0), 0.5% NP-40, and protease inhibitor cocktail (Roche)) supplemented with 2.5% murine RNase inhibitor (New England Biolabs; not included when RNase was used) was added to each pellet. The pellet was homogenized via pipette, transferred to an Eppendorf tube, and incubated at room temperature for 3 min. Lysates were then centrifuged at room temperature for 5 min at 21,000 ×*g*. Lysate supernatants were transferred to a new Eppendorf tube and combined with purified protein to induce LLPS along with any other exogenous reagents used in the experiment. Immediately after mixing, samples were seeded into imaging vessels to observe LLPS. The following imaging vessels were used, including corresponding volumes seeded per well: 20 μl in μ-Slide Angiogenesis with ibiTreat (Ibidi); 125 μl in 8-well lab-Tek chambered coverglass (Nunc); or 20 μl in 384-well glass bottom SensoPlate (Greiner). All experiments that were quantified were performed in 8-well Lab-Tek slides; otherwise, Ibidi slides were used. The 384-plate was only used for the multi-dimensional phase diagram (Fig. 1J). Since temperature is known to influence LLPS of many LCD-containing proteins, all manipulations and analysis following lysis were performed at room temperature (22-25°C).

### Fluorescent microscopy and image analysis

Fluorescent imaging was performed using a Yokogawa CSU W1 spinning disk attached to a Nikon Ti2 eclipse with a Photometrics Prime 95B camera using Nikon Elements software. Imaging was performed through a 60× Plan Apo 1.40NA oil objective and Perfect Focus 2.0 (Nikon) was engaged for all captures. For single timepoint captures using the Ibidi Angiogenesis slides, images were taken at the surface of the sample at 30 min after induction or at the indicated timepoint. Imaging was performed using 488 nm, 555 nm, and 640 nm lasers when applicable along with a capture of DIC. Multipoint time-lapse imaging was performed in Lab-Tek chambered coverglass slides, where 5 xy fields were stored for each condition. Images were taken at each xy position every 10 min. Automated granule detection and measurement was performed using CellProfiler software (Broad Institute) similar to the methods described previously (Mackenzie et al., 2017), excluding association of granules to cells. Briefly, the lysate granules were segmented by applying a Global “minimum cross entropy” approach on the GFP channel. The sizes represent an average of at least 5 separate fields, with each field containing at least 100 segmented granules.

### Multidimensional phase diagram

Lysates at indicated concentrations combined with indicated concentrations of G3BP1 were seeded into a 384-well glass bottom SensoPlate (Greiner) at 20 μl per well. Imaging was performed on a Cytation 5 multi-mode plate reader (BioTek) using a 40× objective and images were captured through transmitted light 60 min after induction.

### Antibody preparation and reagents added exogenously to lysate

Addition of all exogenous reagents occurred simultaneously with the addition of purified G3BP1. Dilutions of reagents were performed in lysis buffer whenever possible. The following antibodies were used: anti-actin (Santa Cruz Biotechnology, sc-1616-R), anti-DCP1a (Abcam, ab47811), anti-FKBP12 (Santa Cruz Biotechnology, sc-28814), anti-TOM20 (Santa Cruz Biotechnology, sc-11415), anti-GOLGA3 (Santa Cruz Biotechnology, sc-292192), anti-caprin1 (Proteintech, 15112-1-AP), anti-PABPC1 (Abcam, ab21060), anti-PRRC2C (Abcam, ab117790), anti-USP10 (Proteintech, 19374-1-AP), anti-UBAP2L (Abcam, ab138309), anti-CSDE (Bethyl Laboratories, A303-158A), anti-DDX21 (Novus, NB100-171855), anti-NPM1 (Sigma, B0556), anti-nucleolin (Abcam, ab22758), anti-TDP43 (Proteintech, 12892-1-AP), Alexa Fluor 647 anti-rabbit (Life Technologies), and Alexa Fluor 647 anti-mouse (Life Technologies), and Alexa Fluor 488 anti-mouse (Life Technologies). For use in lysate LLPS experiments, primary and secondary antibodies were diluted in lysate buffer and combined to pre-conjugate for 1 h at room temperature on an orbital shaker at the lowest speed setting, before being added to lysate alongside purified protein. Total dilutions were 1:1000 for secondary antibodies; primary antibody dilutions were identical to the manufacturers’ recommendation for IF/IHC. Oligo-DT labeled with Cy5 (Genelink, 26-4320-02) and RNase were obtained from Thermo Fisher Scientific (EN0531).

### FRAP

FRAP experiments were performed on a Yokogawa CSU W1 Spinning Disk attached to a Nikon Ti2 eclipse with a Photometrics Prime 95B camera using Nikon Elements software. The light path was split between the port for the spinning disk/acquisition laser and the FRAP lasers, enabling FRAP to occur simultaneously while imaging. All FRAP imaging was taken on a 60× Plan Apo 1.40NA oil objective with Perfect Focus 2.0 (Nikon) engaged. For live-cell FRAP, cells were seeded into 4-well Lab-Tek chambered coverglass (Nunc) with Fluorobrite DMEM media supplemented with 10% FBS and 4 mM L-glutamine. During imaging, cells were maintained at 37°C and supplied with 5% CO_2_ using a Bold Line Cage Incubator (Okolabs) and an objective heater (Bioptechs). To induce stress granules, cells were incubated with 500 μM sodium arsenite (Sigma) for 30 min. Time lapses were acquired as rapidly as possible over the course of 35 s for stress granules and G3BP1-induced granules, and 60 s for nucleoli and NPM1-induced granules, with photobleaching with the 488-nm FRAP laser occurring 2 s into capture. Data were taken from at least n =10 different cells or lysate granules for each condition. In Nikon Elements, ROIs were generated in the photobleached region, a non-photobleached cell, and the background for each time lapse, and the mean intensity of each was extracted. These values were exported into Igor Pro 7.0 (WaveMetrics), where photobleach and background correction were performed, and fit FRAP curves were generated.

### Measurement of protein concentration

To measure the protein content of lysates, Bradford assays were run on lysate supernatants against BSA standards. To measure the total protein content of cells to equalize lysate concentrations in LLPS experiments, 10% of each cell pellet was reserved prior to freezing and lysed in equal volume 2× lysis buffer (ddH_2_O, 100 mM Tris-HCl (pH 7.0), 1% NP-40), and a Bradford assay was run against BSA standards. For LLPS experiments, the lowest concentrated samples were lysed in 225 μl lysis buffer (rather than the standard 250 μl), and the remaining samples were lysed in higher volumes to equalize protein content between samples.

### LC/MS/MS

Lysate granules were induced as described above with either 20 μM G3BP1 or 15 μM NPM1 or a mock purification as a control without addition of purified protein. Granules were allowed to form for 40 min and granules were isolated from the lysate by centrifugation at 2,000 ×g for 5 min. The resulting granules were collected in 2× LDS sample buffer and separated by SDS-PAGE. Proteins were isolated from the gel and digested with trypsin overnight. Samples were loaded on a nanoscale capillary reverse phase C18 column by a HPLC system (Thermo Ultimate 3000) and eluted by a gradient (~90 min). Eluted peptides were ionized by electrospray ionization and detected by an inline mass spectrometer (Thermo Orbitrap Fusion). Database searches were performed using SEQUEST search engine using an in-house SPIDERS software package. MS/MS spectra were filtered by mass accuracy and matching scores to reduce protein false discovery rate to ~1%. The total number of spectral counts for each protein identified was reported by sample. Proteins were included in the granule interactomes if they were found to be enriched at least two-fold in the granule vs. mock samples and had a least two separate peptides identified by LC/MS/MS.

### RNA-seq

Lysate granules were induced with 100 μM G3BP1 and allowed to form for 40 min followed by granule isolation from the lysate by centrifugation at 2,000 ×g for 5 min. RNA was isolated from either the lysate immediately prior to induction with G3BP1 or from the isolated granules. Stranded total RNA was used for RNA-seq analysis in triplicate samples from RNA isolated from both lysate and granules as described below. RNA was quantified using the Quant-iT RiboGreen assay (Life Technologies) and quality checked by 2100 Bioanalyzer RNA 6000 Nano assay (Agilent Technologies), 4200 TapeStation High Sensitivity RNA ScreenTape assay (Agilent Technologies), or LabChip RNA Pico Sensitivity assay (PerkinElmer) prior to library generation. Libraries were prepared from total RNA with the TruSeq Stranded Total RNA Library Prep Kit according to the manufacturer’s instructions (Illumina, PN 20020599). Libraries were analyzed for insert size distribution on a 2100 BioAnalyzer High Sensitivity kit (Agilent Technologies), 4200 TapeStation D1000 ScreenTape assay (Agilent Technologies), or Caliper LabChip GX DNA High Sensitivity Reagent Kit (PerkinElmer). Libraries were quantified using the Quant-iT PicoGreen ds DNA assay (Life Technologies) or low pass sequencing with a MiSeq nano kit (Illumina). Paired end 100 cycle sequencing was performed on a NovaSeq 6000 (Illumina).

### Online supplemental material

Videos 1 and 2 show formation, growth, and fusion of G3BP1-induced lysate granules at low and high power, respectively (related to Fig. 1). Video 3 shows dynamics of G3BP1-induced lysate granules as assayed by FRAP (related to Fig. 1). Videos 4 and 5 show conserved dynamics of cellular stress granules and G3BP1-induced lysate granules as assayed by FRAP (related to Fig. 2). Video 6 shows conserved dynamics of cellular nucleoli and NPM1-induced lysate granules as assayed by FRAP (related to Fig. 4). In all cases, videos of lysate granules are focused on condensates that have settled on the slide surface; videos of granules inside intact cells are focused on cells adhering to the surface of the imaging chamber.

## Acknowledgements

We thank Natalia Nedelsky for editorial assistance. We thank the Center for Advanced Genome Engineering at St. Jude Children’s Research Hospital for assistance with CRISPR-Cas9 modified cell lines, the Center for Proteomics and Metabolomics at St. Jude Children’s Research Hospital for assistance with mass spectrometry analyses, and the Hartwell Center Genome Sequencing Facility for assistance with RNA sequencing. We thank Richard Kriwacki for sharing purified NPM1 and Jessica Hughes for assistance with protein preparation. This work was supported by grants from the Howard Hughes Medical Institute, NIH grant R35NS097974, the St. Jude Research Collaborative on Membraneless Organelles, and the ALS Association (18-IIA-419) to J.P. Taylor. The content is solely the responsibility of the authors and does not necessarily represent the official views of the National Institutes of Health. J.P. Taylor is a consultant for Nido Biosciences and Faze Medicines. The authors have no additional competing financial interests.

## Author contributions

J.P. Taylor conceived and supervised the project. B. Freibaum, J. Messing, and P. Yang designed and/or performed the experiments. B. Freibaum, J. Messing, H.J. Kim, and J.P. Taylor analyzed data and wrote the manuscript.

## Abbreviations

ALS: Amyotrophic lateral sclerosis
DIC: Differential interference contrast
FTD: frontotemporal dementia
LLPS: Liquid-liquid phase separation
nsP3: non-structural protein 3

## Supplemental Information

### Tables

**Table S1. Protein composition of G3BP1-induced lysate granules**. Lysate from a confluent 10-cm plate of U2OS cells was induced with 20 μM G3BP1 purified protein and lysate granules were allowed to grow for 40 min. The resultant granules were captured by centrifugation at 2000 ×*g* for 5 min and subjected to spectral counting on a mass spectrometer. Uninduced lysates were used as a negative control. Spectral counts of 0 were edited to 0.1 within the table to allow for calculation of granule enrichment.

**Table S2. RNA composition of G3BP1-induced lysate granules and stress granules (Khong et al**., **2017)**. Lysate from triplicate confluent 10-cm plates of U2OS cells were collected. RNA from half the lysate was used for RNA-seq analysis while the other half was induced with 100 μM G3BP1 purified protein and lysate granules were allowed to grow for 40 min. The resultant granules were captured by centrifugation at 2000 ×*g* for 5 min. RNA from the granules was purified and subjected to RNA-seq analysis. Comparative analysis was performed with RNA purified from stress granules in U2OS cells (Khong et al., 2017). Raw read counts for both analyses are presented as FPKM and TPM.

**Table S3. Percent composition of RNA by subtype of lysate granules and stress granules (Khong et al**., **2017)**. Percent composition of RNA by subtype of granules formed in G3BP1 lysate granules vs. RNA found in stress granules (Khong et al., 2017).

**Table S4. Protein composition of NPM1-induced lysate granules**. Lysate from a confluent 10-cm plate of U2OS cells was induced with 15 μM NPM1 purified protein and lysate granules were allowed to grow for 40 min. The resultant granules were captured by centrifugation at 2000 ×*g* for 5 min and subjected to spectral counting on a mass spectrometer. Uninduced lysates were used as a negative control. Spectral counts of 0 were edited to 0.1 within the table to allow for calculation of granule enrichment.

### Videos

**Video 1. Formation, growth, and fusion of granules from G3BP1-GFP U2OS lysate induced with purified G3BP1**. Video shows a zoomed-out field of the granule shown in Figure 1F. Imaging was performed using time-lapse spinning disk confocal microscopy of granules settled at the surface of the imaging chamber 5 min after induction of granules with 20 μM purified G3BP1. DIC and G3BP1-GFP images were captured every 0.083 min for 25 min total. Frames per second (FPS) = 30.

**Video 2. Fusion of granules from G3BP1-GFP U2OS lysate induced with purified G3BP1**. The same granule is shown in Figure 1F; also a zoom-in of Video 1. Imaging was performed using time-lapse spinning disk confocal microscopy of granules settled at the surface of the imaging chamber 5 min after induction of granules with 20 μM purified G3BP1. DIC and G3BP1-GFP images were captured every 0.083 min for 25 min total. FPS = 30.

**Video 3. FRAP of G3BP1-GFP from lysate granules induced at various G3BP1 concentrations**. Video corresponds to the FRAP curves shown in Figure 1G. Imaging was performed using time-lapse spinning disk confocal microscopy of granules settled at the surface of the imaging chamber 30 min after induction of granules with purified G3BP1. Images were captured every 0.2 seconds for 35 seconds total. Photobleaching with 488-nm laser occurred 2 seconds into capture. FPS = 15.

**Video 4. FRAP of G3BP1-GFP and GFP-ATXN2L within cellular stress granules**. Video corresponds to stills and FRAP curves shown in Figure 2F-G. Stable G3BP1-GFP or KI GFP-ATXN2L U2OS cells were exposed to 500 μM sodium arsenite for 30 min. Imaging was performed using time-lapse spinning disk confocal microscopy of cells adhering to the surface of the imaging chamber. Images were captured every 0.1 seconds for 35 seconds total. Photobleaching with 488-nm laser occurred 2 seconds into capture. FPS = 30.

**Video 5. FRAP of G3BP1-GFP and GFP-ATXN2L within G3BP1-induced lysate granules**. Video corresponds to stills and FRAP curve shown in Figure 2H-I. Lysates from stable G3BP1-GFP or KI GFP-ATXN2L U2OS cells were induced with 20 μM purified G3BP1. Imaging was performed using time-lapse spinning disk confocal microscopy of granules settled at the surface of the imaging chamber. Images were captured every 0.2 seconds for 35 seconds total. Photobleaching with 488-nm laser occurred 2 seconds into capture. FPS = 15.

**Video 6. FRAP of transfected GFP-NPM1 in the nucleolus and NPM1-induced lysate granules**. Video corresponds to stills and FRAP curves shown in Figure 4B-C. U2OS cells were transiently transfected with GFP-NPM1. Intact transfected cells were imaged to monitor mobility of GFP-NPM1 in the nucleolus. Lysates prepared from transfected cells were induced with 15 μM purified NPM1. Imaging was performed using time-lapse spinning disk confocal microscopy of granules settled at the surface or cells adhering to the surface of the imaging chamber. Images were captured every 0.2 seconds for 60 seconds total. Photobleaching with 488-nm laser occurred 2 seconds into capture. FPS = 30.

